# lncRNA Neat1 drives neuronal histone methylation and age-related memory impairments

**DOI:** 10.1101/531707

**Authors:** Anderson A. Butler, Daniel R. Johnston, Simranjit Kaur, Farah D. Lubin

## Abstract

Histone methylation is critical for the formation and maintenance of long-term memories. Long noncoding RNAs (lncRNAs) are regulators of histone methyltransferases and other chromatin modifying enzymes (CMEs). We investigated how lncRNA *Neat1*-mediated histone methylation contributes to hippocampus-dependent long-term memory formation, using a combination of transcriptomics, RNA binding protein immunoprecipitation, CRISPR mediated gene activation, and behavioral approaches. Suppression of the lncRNA *Neat1* revealed widespread changes in gene transcription as well as perturbations of histone 3 lysine 9 dimethylation (H3K9me2), a repressive histone modification mark that is dysregulated in the aging hippocampus. We identified a Neat1-dependent mechanism of transcriptional repression via H3K9me2 at the *c-Fos* promoter corresponding with observed changes in *c-Fos* mRNA levels. Overexpression of hippocampal *Neat1* via CRISPRa is sufficient to impair memory formation in young adults, recapitulating observed memory deficits in old adults, while *Neat1* suppression in both young and old adult mice improves memory. These results suggest that lncRNA *Neat1* is a potent epigenetic regulator of hippocampus-dependent long-term memory formation.

## INTRODUCTION

While recent efforts have characterized thousands of lncRNAs in the human and mammalian genome, few lncRNAs are as well-studied, as the human Nuclear-Enriched Abundant Transcript 1 (*NEAT1*). *NEAT1* is evolutionarily conserved between rodents and humans, particularly within the 5’ region of the transcript^1^. Multiple isoforms of *NEAT1* exist in rodents and in humans, with the longer of the major isoforms proving essential for phase separation and induction of nuclear paraspeckle assembly^2,3^, while the shorter *NEAT1* transcripts do not appear to be a major regulator of paraspeckle formation^4^. Recent studies have characterized a number of molecular pathways by which *NEAT1* regulates the epigenome, including both paraspeckle-dependent sequestration of transcription factors, as well as paraspeckle-independent roles for *NEAT1* in transcriptional regulation via scaffolding of CMEs^5–7^. Additionally, *NEAT1* itself has been observed to bind numerous genomic loci and to effect regulation of transcription^8,9^.

Research on the human *NEAT1* has been largely focused on its role as an oncogene in various cancers (as reviewed previously^10^), which occurs largely through its regulation of epigenetic mechanisms. However, the rodent homolog *Neat1* is also upregulated in the hippocampus of aging mice^11^ and has recently been linked to multiple cognitive and neurodegenerative disorders, including schizophrenia^12^, Huntington’s Disease^13^, Parkinson’s Disease^14,15^, Alzheimer’s Disease^16^, and epilepsy^17,18^. Furthermore, recent evidence suggests that *Neat1* may play a role in neuroplasticity^17^; however, despite such extensive health relevance, the role of *Neat1* in epigenetic regulation of genes within hippocampal neurons, particularly during long-term memory formation. We used RNA-sequencing, CRISPR mediated gene activation (CRISPRa), and memory tests to investigate the functional role of lncRNA *Neat1* in gene expression dynamics and the role that age-related changes in *Neat1* expression might play in memory deficits in older adults.

## RESULTS

### Expression of the long noncoding RNA NEAT1 is restricted in human CNS tissues

Expression of *NEAT1* is abundant in many cultured cell lines including those characterized in the ENCODE project^19^ (Fig. 1A). However, we observed that in contrast to the abundant expression of *NEAT1* observed in most tissues, the human central nervous system (CNS) as a whole, as well as the hippocampus (outlined in red, Figs. 1B-C) express minimal quantities of *NEAT1*^*20,21*^. Unsupervised hierarchical clustering based on tissue expression of *NEAT1* supports this observation, as CNS tissues segregate cleanly when sorted based on *NEAT1* transcript expression (Fig. 1D, Supplemental Figs. 1A-B).

**Figure 1.**
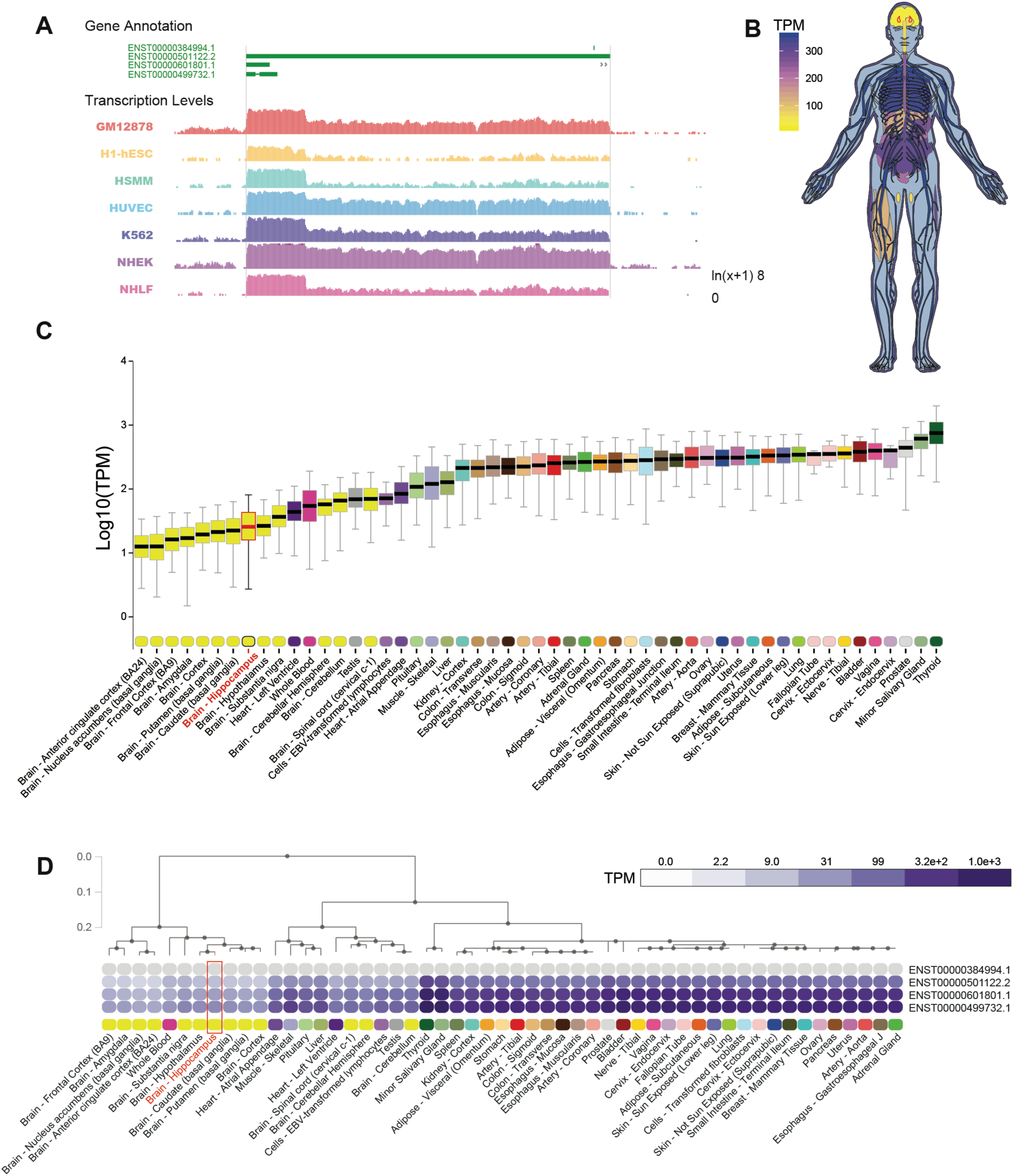
Restricted expression of lncRNA *NEAT1* in human CNS tissues. (A) University of California, Santa Cruz (UCSC) Genome browser track export showing expression of *NEAT1* in seven cell types from ENCODE. **(B)** Human body plot illustrating the expression of *NEAT1* in 53 human tissues from the GTEx project, values shown are the median transcripts per million (TPM) values by tissue, hippocampus outlined in red. **(C)** Bar plots showing median, upper quartile, and lower quartile expression of the *NEAT1* gene (ENSG00000245532.4) in 53 human tissues from the GTEx project; hippocampal expression outlined in red. **(D)** Hierarchical clustering of *NEAT1* based on transcript isoform level expression in 53 human tissues from the GTEx project. Dendrogram scale shows cluster distance. Expression values displayed in the heatmap are the median expression values in TPM for each isoform in each tissue.

Examination of single-cell RNA-seq data from resected human CNS tissue and glioblastoma^22^ further suggests that expression of *NEAT1* within CNS cells is restricted in neurons, while other cell types including astrocytes, oligodendrocytes, and vascular cells express *NEAT1* at higher levels. (Supplemental Figs. 1C-D). This is in contrast to the neighboring lncRNA transcript MALAT1 which appears to be ubiquitously expressed at high levels in all CNS cell types (Supplemental Fig. 1E). Given the growing body of l i terature that has noted overexpression of *NEAT1* in the aging brain^23,24^, as well as the established role of *NEAT1* as a regulator of epigenetic mechanisms, and the recently-described role of *NEAT1* in cognitive disorders such as schizophrenia^12^, we sought to further investigate the role of the lncRNA *NEAT1* on the neuroepigenetic mechanisms of cognition.

### NEAT1 regulates the immediate early gene c-Fos involved in synaptic plasticity

To investigate the role of *NEAT1* at the transcriptomic level, we analyzed a publicly available RNA-seq dataset from iPSC-derived human neurons. Antisense oligo (ASO) knockdown of *NEAT1* in KCl-treated human neurons revealed an extensive cohort of differentially expressed mRNAs. Knockdown alone was not sufficient to perturb the transcriptome in resting iPSC-derived human neurons, as evidenced by an imperfect separation via unsupervised hierarchical clustering prior to KCl stimulation (Fig. 2A). In contrast, *NEAT1*-knockdown appears to dramatically potentiate KCl-driven differential expression of many genes (Fig. 2A-B).

**Figure 2.**
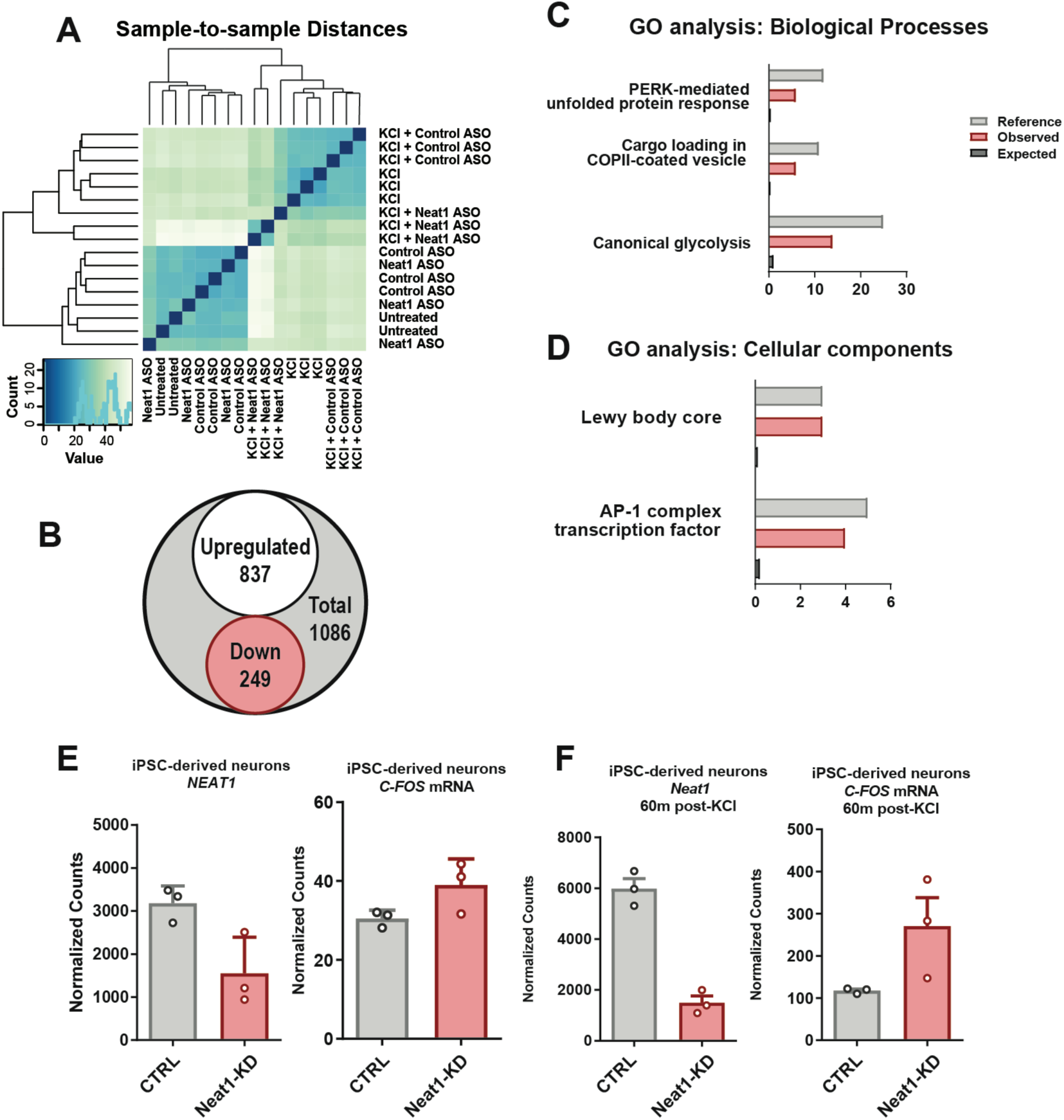
*NEAT1* regulates expression of *C-FOS* mRNA and the AP-1 complex in iPSC-derived human neurons. **(A)** Unsupervised hierarchical clustering transcriptomes from *Neat1* and KCl treated iPSC-derived human neurons, based on DESeq2-normalized counts **(B)** Venn diagram depicting the total number of differentially expressed genes detected between KCl+ Control_ antisense oligonucleotide (ASO) and KCl+*Neat1*_ASO groups via DESeq2. **(C-D)** Gene Ontology (GO) term enrichment for DE genes depicted in panel B. All GO terms shown showed statistically significant enrichment (BH corrected *p* < 0.05) **(E-F)** Normalized count values for lncRNA *NEAT1* and *C-FOS* mRNA either prior to **(E)** or after **(F)** KCl treatment of iPSC-derived neurons.

To gain some insight into the health relevance for observed *NEAT1*-mediated changes in gene expression in human neurons, we queried the annotated disease classes from the Genetic Association Database via the Database for Annotation, Visualization and Integrated Discovery (DAVID)^25^, and observed significant enrichment for three disease classes: cancer, renal, and aging. (Supplemental File 2). *NEAT1*-regulated genes appear to be non-randomly distributed among annotated biological processes (Fig. 2C), molecular functions (Supplemental File 2) and cellular components (Fig. 2D). Significant gene ontology (GO) term enrichment was partially consistent with previous observations of the *NEAT1* regulatory axis, as we observed significant regulation of GO terms associated with viral gene expression; however, we also observed significant enrichment of GO terms important for hippocampal function, including the transcription factor AP-1 complex (GO:0035976, Fig. 2C-D).

The human Fos proto-oncogene (*FOS*, also known as *C-FOS*), a critical component of the AP-1 transcription factor subunit appeared to be overexpressed in human neurons after knocking down *NEAT1* both in quiescent and KCl-stimulated neurons and has a known relevance for hippocampus-dependent memory formation^26^ (Figs. 2E-F). Thus, we selected the murine homolog *c-Fos* as a candidate gene for further studies of *Neat1*’s regulatory potential.

### Neat1 is regulated by neuronal excitability and controls c-Fos gene expression

As modeling *Neat1* expression changes in response to *in vivo* neuronal activity and behavioral experience required the mammalian model organisms, we next sought to examine the regulatory capacity of *Neat1* in rodent neurons. For this purpose, we knocked down murine *Neat1* in the mouse Neuro-2a (N2a) cell line using small interfering RNAs (siRNAs). We observed that 24h after treatment with *Neat1*-targeting siRNAs (Fig. 3A), expression of the *c-Fos* mRNA was significantly upregulated (Fig. 3B).

**Figure 3.**
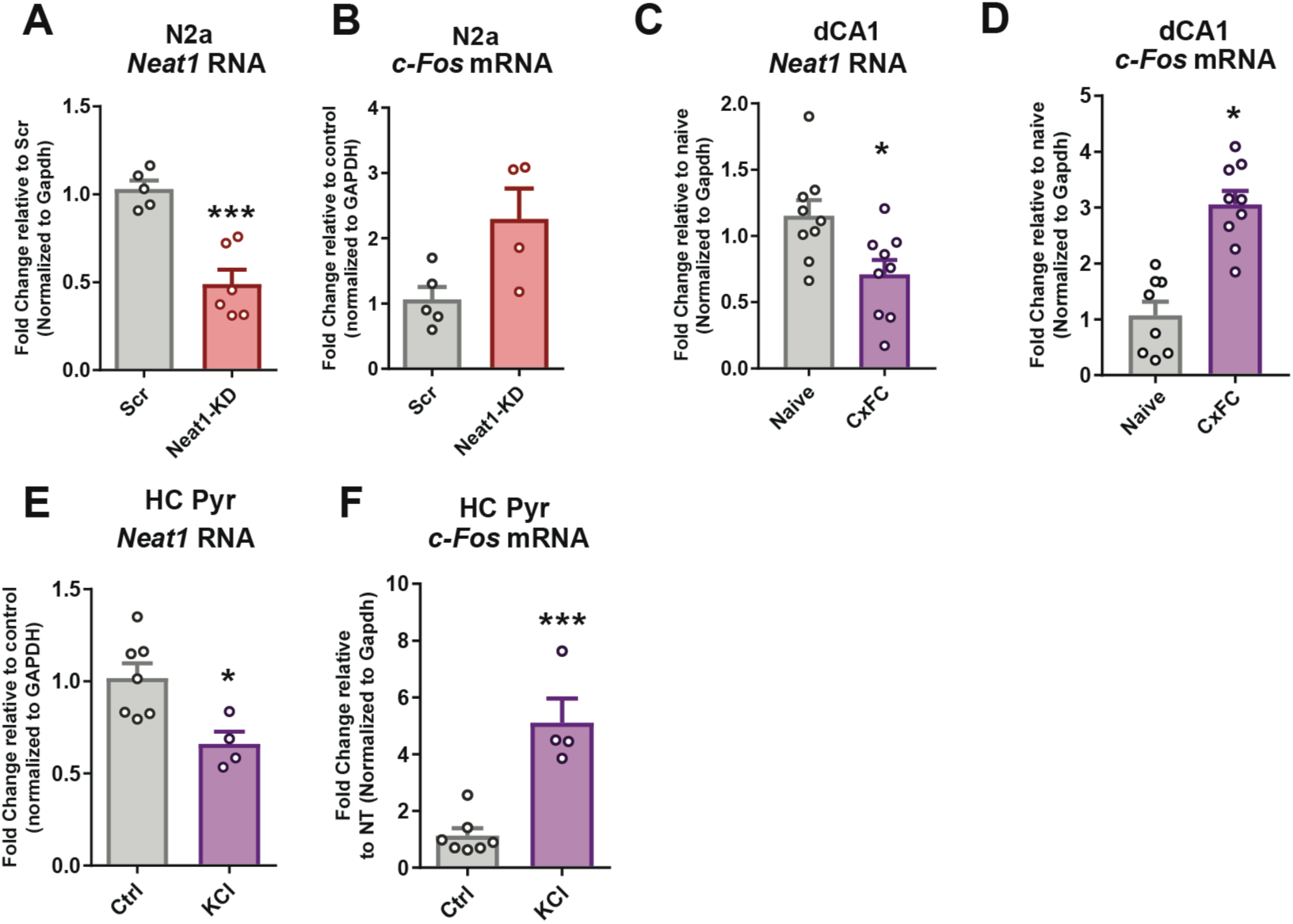
*Neat1* regulates expression of *c-Fos* mRNA in murine neuronal cells. **(A)** siRNA treated murine N2a cells show significantly reduced abundance of *Neat1* transcript. (*n* = 5,6; *p* < 0.0005) **(B)** Expression of *c-Fos* mRNA after treatment with *Neat1*-targeting siRNAs (*n =* 5,4; *p* = 0.0335)**. (C-D)** *Neat1* expression is decreased (**C**; *n* = 9,9; *p* < 0.0148) and *c-Fos* expression is increased (**D**; *n* = 8,9; *p* < 0.0001) *in vivo* in dorsal CA1 1h after training in contextual fear conditioning **(E-F)** Depolarization of rodent primary pyramidal neurons with KCl is sufficient to significantly reduce expression of *Neat1* **(E;** *n =* 7,4; *p* = 0.0147) and reproduce commonly observed increases in *c-Fos* transcription **(F**; *n =* 7,4; *p* = 0.0003).

Interestingly, while our observations of c-*Fos* transcript expression in murine neurons recapitulated observations from human neurons, we observed that expression of the immediate early genes Egr1 and Btg2 were not overexpressed in mouse (Supplemental Fig. 2A-B) as they were in human neurons (Supplemental File 2), suggesting that there are species-specific regulatory differences in the *Neat1* regulatory axis.

As mice with the *c-Fos* gene knocked out in the CNS show a specific loss of hippocampus-dependent spatial and associative learning tasks^26^, we next sought to investigate the relevance of *Neat1* expression during memory consolidation after a hippocampus-dependent learning task. One hour after training in contextual fear conditioning we observed a significant reduction in the expression of *Neat1* in the dorsal hippocampus coinciding with previously reported increases in expression of the *c-Fos* mRNA (Figs. 3C-D). As baseline expression of *Neat1* in neurons is expected to be quite restricted compared to other cell types, we next stimulated neurons with KCl to ascertain the effect of activity on the expression of *Neat1*. Consistent with recent reports^17^, we observed that KCl stimulation drives a rapid reduction in *Neat1* expression in both N2a cells (Supplementary Fig. 2E) and primary hippocampal pyramidal neurons, as recently reported (Figs. 3E-F), and consistent with the effects of context fear conditioning *in vivo*.

### Neat1 regulates H3K9me2 globally and controls c-Fos promoter H3K9me2 and gene expression

We next sought to investigate the *c-Fos*-relevant mechanisms of *Neat1*-orchestrated transcriptional control. To accomplish this, we used publicly available data assaying *Neat1* chromatin binding via capture hybridization analysis of RNA targets and high-throughput sequencing (CHART-seq) in human MCF7 cells^8^. After mapping *Neat1*-bound peaks to the nearest transcription start sites, we observed that only a small subset of genes directly bound by *Neat1* are differentially expressed either after *Neat1* knockdown or in the context of neuronal activation. However, we observed significant enrichment of *Neat1* binding near genes associated with histone methyltransferase activity, including the H3K9 dimethyltransferase Ehmt1 (also known as GLP) (Figs. 4A-B; Supplemental File 3).

**Figure 4.**
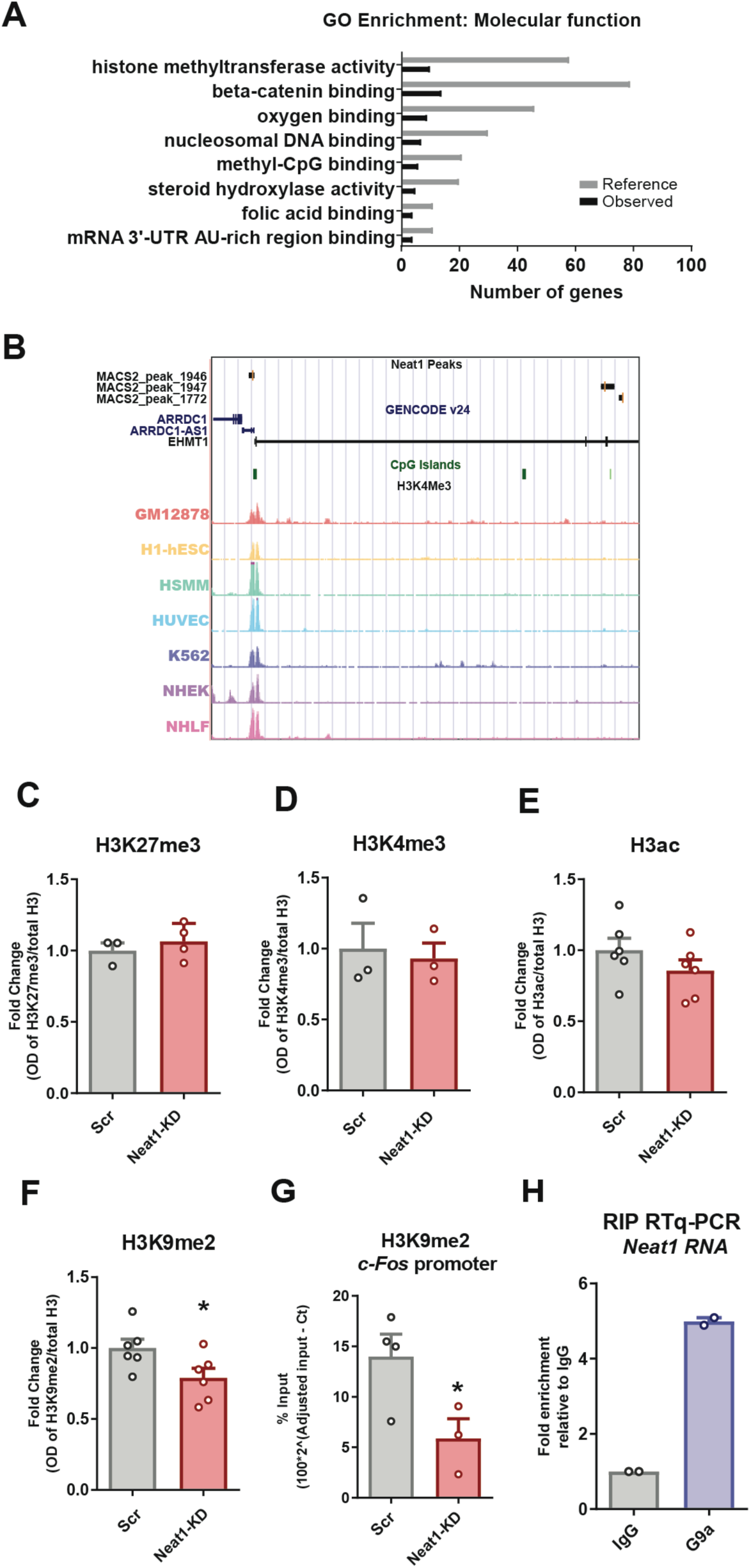
*Neat1* modulates neuronal H3K9me2. **(A)** *NEAT1* CHART-seq peaks were mapped to the nearest gene transcription start site, and functional enrichment was assessed using ChIP-ENRICH, with histone methyltransferase activity being noted as a significantly enriched GO term (BH corrected *p* < 0.05 for all terms shown). **(B)** UCSC genome browser plot showing *NEAT1*-binding peaks overlapping the human *EHMT1* gene. **(C-F)** Graphs depicting changes in histone modifications in N2a cells after siRNA knockdown of *Neat1* **(C)** H3K27me3 (n = 3; p = 0.4716) **(D)**H3K4me3 (*n* = 3; *p* = 0.7548) **(E)**H3ac (n = 6; p = 0.2377) **(F)** H3K9me2 (*n* = 6; *p* = 0.0456) **(G)** ChIP-qPCR assay indicating a loss of H3K9me2 at the *c-Fos* gene promoter in N2a cells (n = 4,3; p = 0. 0 4 7 2) **(H)** RNA binding protein immunoprecipitation for Ehmt2/*Neat1* interaction (n = 2)

As *c-Fos* has previously been observed to be regulated by the Ehmt1/2 complex in the context of hippocampus-dependent memory formation^27^, we next sought to investigate the role of *Neat1* in the regulation of histone methylation and H3K9me2 specifically. After knockdown of *Neat1* in neuronal cells, we observed that H3K9me2 is reduced at a global scale while the expression of other histone modifications are unchanged (Fig. 4C-F). To ascertain whether the lncRNA *Neat1* physically associates with the H3K9me2 methyltransferase complex in neurons, we performed RNA binding protein immunoprecipitations against the Ehmt2 subunit of the obligatory Ehmt1/2 heterodimer^27–29^. Consistent with recently published results^6^, we observed interaction between *Neat1* and the H3K9me2 methyltransferase Ehmt2, suggesting multiple possible modes of action for *Neat1*-mediated regulation of H3K9me2.

To assess the functional relevance of the *Neat1*-H3K9me2 regulatory axis on the expression of *c-Fos* mRNA, we performed ChIP in conjunction with qPCR at the *c-Fos* promoter. We observed that after *Neat1* knockdown with siRNAs, H3K9me2 at the *c-Fos* promoter was significantly depleted (Fig. 4G), consistent with observed changes in gene expression (Fig. 3B), while H3K9me2 within the *c-Fos* gene body were not significantly changed (Supplementary Fig. 2F).

### Neat1 knockdown regulates hippocampal memory formation and the epigenetic landscape at the c-Fos promoter in vivo

Having demonstrated that *Neat1* represses the epigenetic landscape and neuronal expression of the memory-critical *c-Fos* gene, we next sought to investigate the functional role of *Neat1* expression on *c-Fos* promoter methylation and memory formation *in vivo*.

To ask whether *Neat1* expression impacts hippocampus-dependent memory formation, we knocked down expression of *Neat1* in hippocampal area CA1 by directly infusing *Neat1*-targeting siRNAs or non-targeted siRNAs and assayed long term memory function using contextual fear conditioning, a hippocampus-dependent memory task (Fig. 5A). We observed that five days after intra-CA1 injection of *Neat1*-targeting siRNAs, a time when we observe significant reduction in expression of *Neat1* (Supplemental Fig. 3A), mice had no differences in freezing behavior during the training phase of contextual fear conditioning, either before or after delivery of the foot shock (Fig. 5B). However, when returned to the training context 24h later, mice that received *Neat1*-targeting siRNAs displayed significant increases in freezing behavior relative to mice that received non-targeting siRNAs (Fig. 5C).

**Figure 5.**
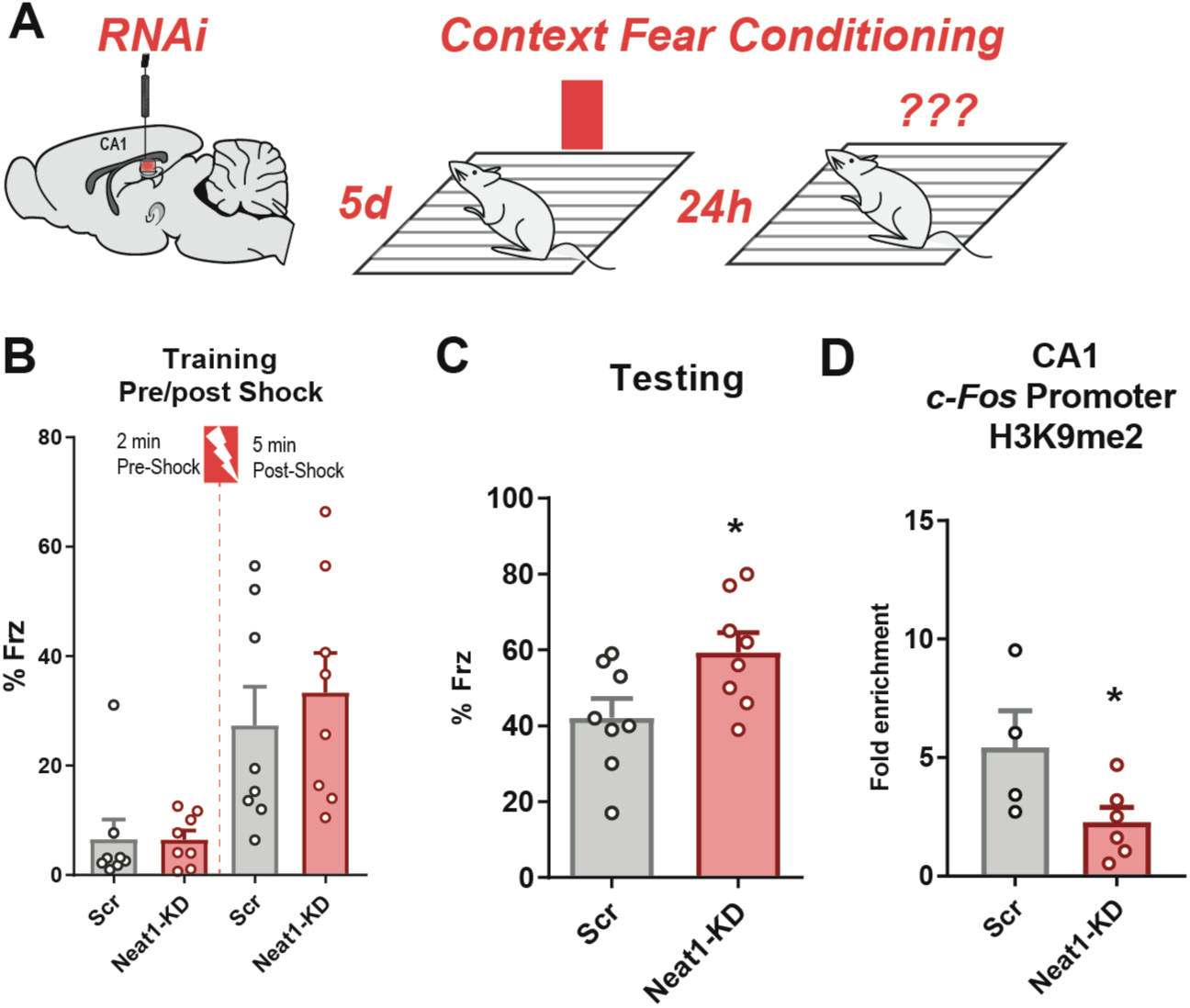
*Neat1* knockdown regulates *c-Fos* promoter methylation *in vivo* and improves long-term memory. **(A)** Graphic depiction of siRNA infusion into hippocampal area CA1 and single-pairing contextual fear conditioning paradigm. Briefly, male C57BL/6 mice were trained 5d after bilateral infusion of siRNAs and tested 24h after training. **(B)** Freezing behavior as a percent of epoch during training phases of the contextual fear conditioning paradigm. No significant difference detected for either the Pre-shock (*n* = 8; *p* = 0.9826) or Post-shock (*n* = 8; *p* = 0.5626) epochs. **(C)** Freezing behavior as a percent of total time during the 5-min test trial (*n* = 8; *p* = 0.0307). **(D)** *c-Fos* promoter H3K9me2 remained depleted after *Neat1*-knockdown 5d after the conclusion of behaviour experiments (*n* = 18,18; *p* = 0.0450)

To determine whether *Neat1* expression impacts *c-Fos* promoter methylation *in vivo*, we sacrificed an additional cohort of behaviorally naïve animals five days after injection of siRNAs and performed ChIP-qPCR assays on one hemisphere of dorsal CA1 tissue collected from around the injection site. Consistent with our results in cultured neurons (Fig. 4G), we observed that concurrently with *Neat1* knockdown five days after infusion with siRNAs, *Neat1* knockdown significantly reduced H3K9me2 at the *c-Fos* promoter in dorsal area CA1 (Fig. 5E). Thus, we hypothesized a model in which *Neat1* expression might be regulating memory formation via epigenetic repression of *c-Fos*.

### Mimicking age-related upregulation of Neat1 is sufficient to respectively restore or impair hippocampus-dependent memory formation

Numerous studies have reported overexpression of *Neat1* in senescing cells, as well as in aging CNS tissues in both humans and mice^11,30^. Upon comparing publicly available hippocampus RNA-seq datasets from 3 month-old young mice versus 24 month-old aged mice we observed upregulation of *Neat1* relative to young animals, consistent with previously reported results^11^ (Fig. 6A), as well as downregulation of *c-Fos* gene expression (Fig. 6B), consistent with previously reported age-associated hippocampus-dependent memory impairments^11^.

**Figure 6.**
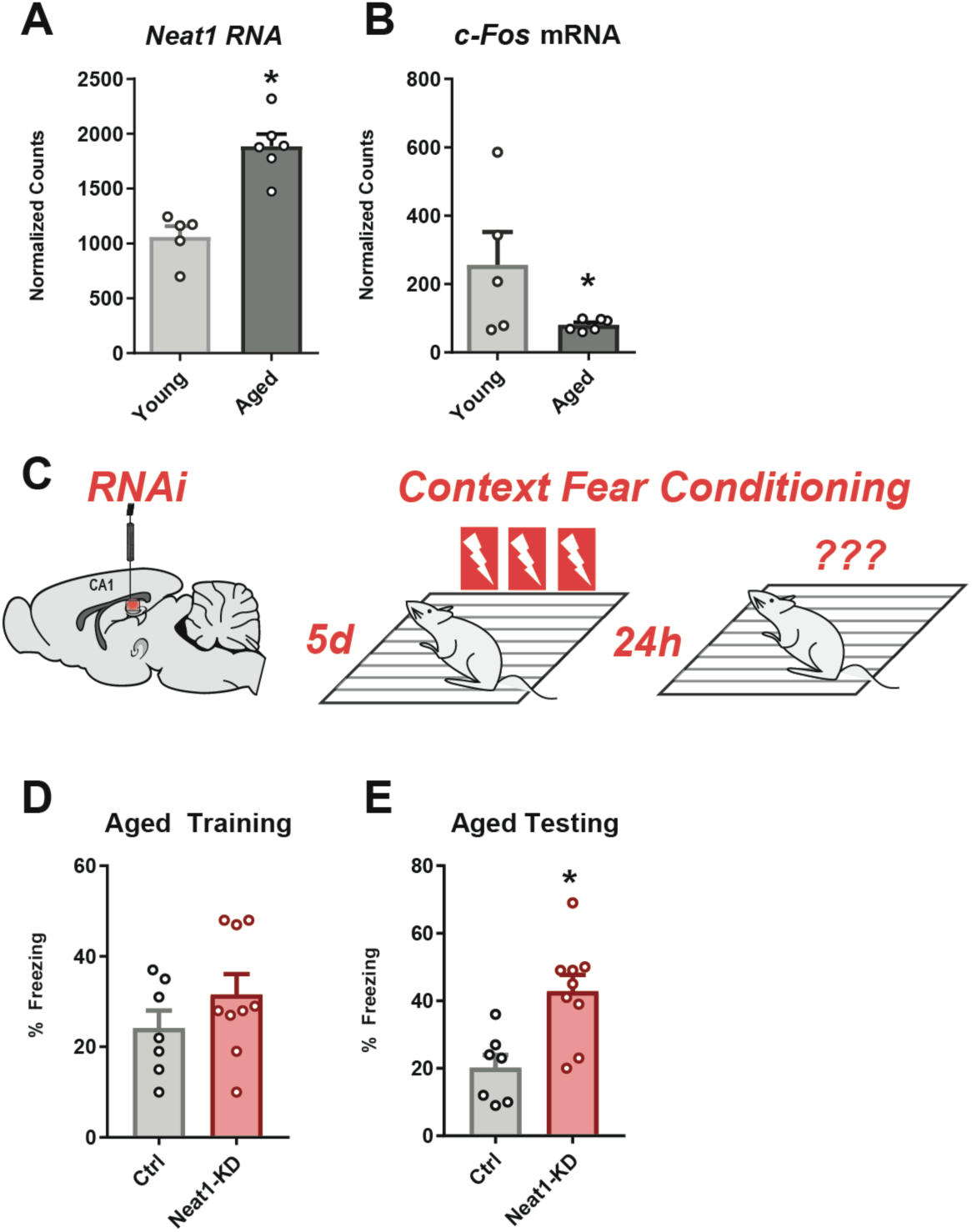
*Neat1*-knockdown improves long-term memory in aged animals. **(A-B)** DESeq2-generated normalized counts of RNAseq data from 3mo and 24mo C57/B6 mice. *Neat1* expression **(A)** was significantly elevated and *c-Fos* mRNA **(B)** was significantly repressed in aged hippocampi relative to the hippocampi of young mice**. (C)** Graphic depiction of siRNA infusion into hippocampal area CA1 and single-pairing contextual fear conditioning paradigm. Briefly, 18mo old male C57/ B6 mice were trained 5d after bilateral infusion of siRNAs and tested 24h after training. **(D-E).** Aged mice (18 month-old) were trained with three pairings of shock with a novel context after knockdown of *Neat1* and demonstrated no significant difference during training **(D)** (*n* = 7,9; *p* = 0.2496), but significantly enhanced freezing 24h after testing **(E)** (*n* = 7,9; *p* = 0.0039).

**Figure 7.**
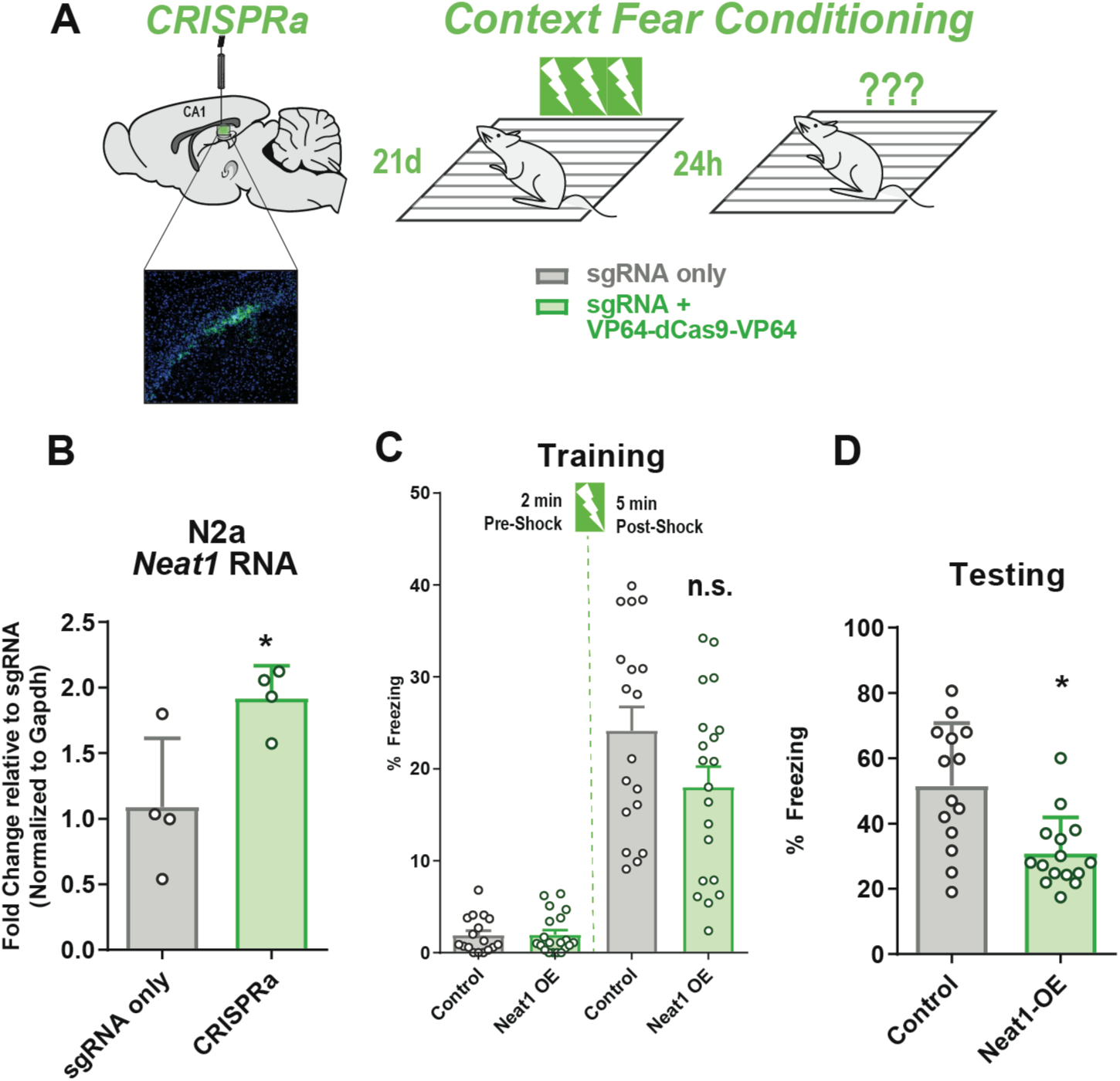
Mimicking age-related *Neat1* overexpression via CRISPRa impairs hippocampal memory formation. **(A)** Graphic depiction CRISPRa system infusion into hippocampal area CA1, with visualization of hippocampal expression of EGFP fluorescent marker, and three-pairing contextual fear conditioning paradigm. Briefly, male C57BL/6 mice (3-7 month-old) were trained 21d after bilateral infusion of either sgRNA plasmid alone or co-delivered with a transcription-activating dCas9-effector protein and tested 24h after training. **(B)** Confirmation of efficacy of CRISPRa system to upregulate *Neat1* expression in murine neurons **(C).** Freezing behavior as a percent of epoch during training phases of the contextual fear conditioning paradigm. No significant difference detected for either the Pre-shock (*n* = 18; *p* = 0.3476) or Post-first shock epochs (*n* = 18; *p* = 0.0665). **(D)** Freezing behavior as a percent of total time during the 5 min test trial (*n* = 18; *p* = 0.0450).

We next tested whether hippocampus-dependent memory formation might be improved in aged mice by knockdown of *Neat1*. To this end, we knocked down expression of *Neat1* in the hippocampal area CA1 of 18-19 month-old mice, an age at which we have previously observed significant upregulation of H3K9me2 in the aging rat hippocampus^31^ (Supplemental Fig. 4), by directly infusing *Neat1*-targeting siRNAs or non-targeted siRNAs and assayed long term memory function using contextual fear conditioning, with three pairings of the shock to the novel context (Fig. 6C). We observed that knockdown of *Neat1* in the dorsal hippocampus of aged mice resulted in significant improvements in freezing after 24h (Figs. 6D-E), but not during training, similar to results seen in young mice (Fig. 5C).

We next sought to test the sufficiency of *Neat1* overexpression to regulate performance in memory tasks, we designed a single guide RNA (sgRNA) targeting *Neat1* for overexpression from the endogenous locus (Fig. 7A, Supplemental Fig. 3B-C), and delivered the CRISPRa system *in vivo* into dorsal CA1 via *in vivo* transfection (Fig. 7B). Mice were then trained in contextual fear conditioning with three pairings of the shock to the novel context (Fig. 6D). Animals overexpressing *Neat1* from the endogenous locus (*Neat1*-OE) had no significant differences in freezing during the training period, either before or after exposure to the unconditioned stimulus (Fig. 6E); however, when returned to the training context 24 h after training, *Neat1*-OE animals froze significantly less than control animals which received only the sgRNA plasmid (Fig. 6F), suggesting that elevated *Neat1* in area CA1 is sufficient to impair hippocampus-dependent memory formation.

## DISCUSSION

While previous studies have observed regulatory roles for the lncRNA *NEAT1*, including that *NEAT1* localizes to chromatin and governs chromatin modification^8,9^, little work has been done to resolve this regulatory role of *NEAT1* in the context of long-term memory formation. RNA sequencing analysis revealed that the human lncRNA *NEAT1* binds to the *EHMT1* locus and that *NEAT1* knockdown regulates neuronal *EHMT1* expression (Supplemental Data S1). We observed that murine *Neat1* acts as a potent regulator of H3K9me2 both in cultured cells and *in vivo* (see Fig. 4). Due to recent observations that *NEAT1* interacts directly with Ehmt2^6^, an observation which we ourselves have reproduced via RIP (see Fig. 4), we cannot yet ascertain whether transcriptional control of *EHMT1* or direct interaction with the repressor complex is the rate-limiting factor for H3K9me2 abundance. This intricate multipoint interaction is perhaps illustrative of the intricate systems of regulatory feedback which are thought to control epigenetic mechanisms. Nonetheless, knockdown of *Neat1* was sufficient to perturb this system and to result in both bulk and site-specific changes in H3K9me2 in neurons.

Previous investigations as to the epigenetic regulatory role of *Neat1* have resulted in paradoxical observations to the effect that *Neat1* binds to genomic loci and mediates activation of transcription^9^, but that suppression of *Neat1* expression results primarily in increased neuronal gene expression^17^. We show here that *Neat1* induces widespread regulation of neuronal H3K9me2, potentially resolving this dilemma and further explaining age-related increases in H3K9me2 previously observed in the hippocampus. Moreover, we observed that *Neat1* expression is correlated with H3K9me2 globally as well as at the promoter of the aging-repressed memory-related gene *c-Fos*. While *Neat1* has been observed to act on and via numerous epigenetic mechanisms, to our knowledge this is the first observation that *Neat1*-mediated epigenetic mechanisms are sufficient to govern cognitive function.

Studies of the neuronal impact of *Neat1* expression have thus far been limited to the context of neurological disorders, and in many cases to cultured neuronal cells. Our observations suggest that *Neat1* plays a regulatory role in neuronal H3K9me2 both in cultured neurons and *in vivo*, and that increases in *Neat1* might play a significant role in the age-related decline of hippocampus-dependent memory formation. In humans, expression of *NEAT1* is generally limited in the CNS, and overexpression is a common hallmark of several neurological disorders. While experimental reduction of *Neat1* has very recently been shown to have therapeutic potential in the context of such disorders, the impact of age-related changes in expression has remained unexplored until now. Interestingly, while our experiments were designed to investigate the age-related impact of *Neat1*, we note that recent experiments have described neuroinflammation-mediated increases in *Neat1* expression^32^, and that the findings described in this manuscript implicate *Neat1* as a potential mechanism by which neuroinflammation might impact memory.

While the experiments described here are largely sufficient to explain prior observations of elevated H3K9me2 in the aging hippocampus (Supplementary Fig. 4;^31^, our experiments indicate that increased expression of *Neat1* is not sufficient to explain all of the aging-related neuroepigenetic changes observed in this region. It is likely that many hippocampal lncRNAs have distinct or overlapping roles in the regulation of the neuroepigenetic aging process. Indeed, human *NEAT1* itself has been observed to associate with multiple chromatin modifying enzymes ^6,33–35^. Although we did not detect significant regulation of histone modifications other than H3K9me2 at the global level after knockdown of *Neat1*, the absence of such observations does not preclude the existence of biologically or behaviorally meaningful epigenetic regulation that is more limited in scope and might be uncovered in future studies with a large-scale sequencing approach.

In this work, we demonstrate that *Neat1* regulates a critical transcriptional pathway for hippocampus-dependent memory in rodent neurons *in vitro, in vivo*, and likewise in iPSC-derived human neurons. While attempts to establish the functionality of the evolutionarily conserved lncRNA *Neat1* have met with limited success, little has yet been done to functionally characterize *Neat1* in the context of cognition. Here, we observed that the lncRNA *Neat1* may serve as an endogenous molecular brake on the formation of hippocampus-dependent spatial memories.

## Acknowledgements

This work was supported by grants from the National Institute of Mental Health (FDL: MH082106, MH097909) and the Evelyn F. McKnight Brain Research Foundation.

## Declaration of Interest

The authors declare no competing financial interests.

## METHODS

### Animal housing

Naïve 3-7 month-old or 18 month-old C57BL/6 mice were group housed (2-7 animals/cage) in plastic cages with *ad libitum* access to food and water and were maintained on a 12-h light/dark cycle. All behavioral tests were conducted during the light cycle, and all procedures were approved by the University of Alabama at Birmingham Institutional Animal Care and Use Committee and done in accordance with the National Institute of Health ethical guidelines.

### Cell culture

N2A cells were maintained in DMEM supplemented with 10% FBS. After thawing, the cells were passaged a minimum of two times prior to use in experiments. The cells were kept at 37°C in a 5% CO_2_ incubator. Dissociated cultures of hippocampal pyramidal cells were obtained from embryonic day 18 rat embryos as described previously ^36^. Briefly, timed-pregnancy female Sprague-Dawley rats were terminally anesthetized and embryos were removed from the uterus, then transferred to Hank’s balanced salt solution (HBSS, Gibco) for dissection. Primary rat hippocampal neurons were dissociated via incubation with papain for 20 min at 37°C, rinsed in HBSS, then resuspended in Neurobasal medium (Gibco) and further mechanically dissociated by passing through a series of progressively smaller fire-polished glass Pasteur pipettes. The resulting suspension was passed through a 70-µm cell strainer and plated on poly-L-lysine coated 24-well plates (∼7.5×10^4^ cells per well). Cells were maintained for 2 weeks in Neurobasal supplemented with B-27 and Glutamax (Thermo Fisher Scientific) at 37°C and 5% CO_2_. For KCl stimulation, 6.25μL 1M KCl (Sigma) was added to two-week in vitro cultures, for a final concentration of 12.5mM KCl.

### siRNA Delivery

Young, 3-7 month-old mice were anesthetized with an intraperitoneal injection of ketamine-dexmedetomidine and received bilateral intra-CA1 injections of Lincode SMARTpool siRNAs (Dharmacon) targeting the murine *Neat1* (#R-160022-00-0005) or a negative control (#D-001320-10-05), conjugated with *in vivo* JetPEI (PolyPlus Transfection), an *in vivo* transfection reagent, at the stereotaxic coordinates (AP -2.0mm, ML ±1.5 mm, DV-1.7 mm) with respect to bregma. Aliquots of siRNA stocks (100μM) were diluted to a concentration of ∼2.5μM and conjugated with *in vivo* JetPEI on the day of surgery. Infusions were given over a 10 min period (0.1μL per min) for a total volume of 1μL per hemisphere. After a 48 h recovery period, mice were handled daily for >3min and trained in contextual fear conditioning at five days post-surgery. Aged (18-19 month old) mice were treated similarly but were anesthetized with vaporized isoflurane (3% induction, 2% maintenance). Mice were sacrificed at ten days post-surgery and dorsal area CA1 was harvested from each hemisphere.

### CRISPRa Delivery

Mice were anesthetized with an intraperitoneal injection of ketamine-dexmedetomidine and received bilateral intra-CA1 injections of a guide RNA expression vector driven by the murine U6 promoter and targeting the murine *Neat1* promoter region (Addgene #44248) either alone or in conjunction with an expression vector coding for the S. pyogenes dCas9 fused to two copies of the VP64 transactivator domain (Addgene #59791). Endotoxin-free plasmids were purified using an endotoxin-free plasmid DNA purification kit (Machery-Nagel) and aliquoted to minimize freeze-thaw cycles. Endotoxin-free plasmid stocks were diluted to a final concentration of ∼500ng/uL in sterile 10% glucose and incubated with in vivo JetPEI for 15 min at room temperature on the day of surgery. The resulting transfection complex was delivered via direct infusion at the stereotaxic coordinates (AP -2.0mm, ML ±1.5 mm, DV-1.4 mm) with respect to bregma. Infusions were given over a 10 min period (0.1μL per min) for a total volume of 1μL (∼500ng plasmid DNA) per hemisphere.

### Contextual fear conditioning

Mice were trained to either a weak or strong contextual fear conditioning (CFC) paradigm in a novel context, and long term memory was assessed upon returning the animals to the training context 24h after training. The weak CFC paradigm consisted of a 118s baseline followed by a single shock (0.5mA, 2 sec) pairing in the novel context, while the strong CFC paradigm consisted of a 119-sec baseline followed by three shock pairings (0.5mA, 1s) with interleaved rest periods of 59 sec each. Twenty-four h after training, animals were placed back into the training context for five min to test retention. Freezing behavior was scored by Med Associates software.

### Collection of whole area CA1

One hour after training, the whole brain was removed by gross dissection and placed in oxygenated (95%/5% O_2_/CO_2_) ice-cold cutting solution (110mM sucrose, 60mM NaCl, 3mM KCl, 1.25mM NaH_2_PO_4_, 28mM NaHCO_3_, 0.5mM CaCl_2_, 7mM MgCl_2_, 5mM glucose, and 0.6mM ascorbate). The CA1 region of the hippocampus was then microdissected from each hemisphere and flash frozen on dry ice.

### Collection of dorsal area CA1

Animals were sacrificed by cervical dislocation after overdosing with isoflurane at experiment-specific time points, and the whole brain was rapidly removed and immediately frozen on dry ice. The CA1 region of the dorsal hippocampus was then dissected out with the aid of a mouse brain matrix (Harvard Apparatus) to collect the area of CA1 targeted by siRNA or CRISPRa infusions. All tissue was stored at -80°C prior to processing.

### Western blotting

Normalized proteins (2-10µg) were separated via electrophoresis on either 10% or 20% polyacrylamide gels, transferred onto an Immobilon-FL membrane using a turbo transfer system (Biorad). Membranes were blocked in Licor blocking buffer and probed with the following primary antibodies for histone H3 (1:1000; Abcam #ab1791), H3K9me2 (1:1000; Millipore #07-441), H3K27me3 (1:1000; Millipore #07-449), H3K4me3 (1:1000; Millipore #04-745). Secondary goat anti-rabbit 700CW antibody (1:20,000; Licor Biosciences) was used for detection of proteins using the Licor Odyssey system. All protein quantification was done using ImageStudio Lite software (Licor).

### Reverse transcription qPCR (RT-qPCR)

Section text RNA was extracted from isolated CA1 or cultured cells using Trizol reagent according to the manufacturer’s recommended protocol (Fisher). RNA yield was quantified spectrophotometrically (Nanodrop 2000c), and ∼200ng of RNA was DNAse treated (Amplification grade DNAse I, Sigma), converted to cDNA (iScript cDNA synthesis kit; Biorad), and PCR amplified on the CFX1000 real-time PCR system (BioRad), with primer annealing temperatures of 60°C. See supplemental table for full descriptions of primers used. All data were analyzed using the delta delta Ct method.

### Cell culture

ChIP was performed as described previously^31,37^. Briefly, samples were fixed in PBS with 1% formaldehyde for ten minutes at room temperature, chromatin was sheared using a Bioruptor XL on high power, lysates cleared by centrifugation and diluted in TE and RIPA buffer. Extracts were mixed with MagnaChIP protein A/G beads and immunoprecipitations were carried out at 4°C overnight with 5µg primary antibody (anti-H, Abcam #ab40542; anti-Ezh2, #ab3748) or no antibody (control). Immune complexes were sequentially washed with low salt buffer (20 mM Tris, pH 8.0, 0.1% SDS, 1% Triton X-100, 2 mM EDTA, 150 mM NaCl), high salt buffer (20 mM Tris, pH 8.1, 0.1% SDS, 1% Triton X-100, 500 mM NaCl, 1 mM EDTA), LiCl immune complex buffer (0.25 M LiCl, 10 mM Tris, pH 8.1, 1% deoxycholic acid, 1% IGEPAL-CA630, 500 mM NaCl, 2 mM EDTA), and TE buffer, and eluted into 1xTE containing 1% SDS. Protein-DNA crosslinks were reversed by heating at 65°C overnight. After proteinase K digestion (100µg; 2h at 37°C), DNA was purified by phenol/chloroform/isoamyl alcohol extraction and ethanol precipitation. Immunoprecipitated DNA was quantified via spectrophotometry (Nanodrop 2000c) and ∼15ng of DNA from each sample was assayed via quantitative real-time PCR using primers specific to mouse genes of interest. See supplemental table for full descriptions of primers used.

### RNA binding protein immunoprecipitation (RIP)

RIP was performed as described previously^38^. Briefly, ∼5ug of primary antibody against Ehmt2 (Abcam #ab40542), Ezh2(Abcam #ab3748) or normal rabbit IgG (Cell signaling) were conjugated with 25uL MagnaChIP protein A/G beads (EMD Millipore). Freshly harvested nuclear pellets from at least 10^6 N2a cells were sheared via Dounce homogenization (15-20 strokes) in RIP buffer (150 mM KCl, 25 mM Tris pH 7.4, 5 mM EDTA, 0.5 mM DTT, 0.5% NP40, 1x Protease Inhibitor Cocktail (Sigma), 100 U/ml SUPERASin (Ambion), cleared via centrifugation at 13,000 RPM to remove nuclear membrane and debris, and split into fractions for IP. Sheared nuclear extracts were mixed with antibody conjugated MagnaChIP protein A/G beads and immunoprecipitations were carried out at 4°C for four hours. Beads were then immobilized on a magnetic tube rack, and immune complexes were sequentially washed three times with RIP buffer. Beads were then resuspended in 1mL of Trizol (Thermo Fisher), and coprecipitated RNAs were isolated according to the manufacturer’s recommended protocol. RT-qPCR for *Neat1* was then performed as described above.

### Statistical analyses

Data from all experiments were analyzed using Analysis of Variance (ANOVA) with Fisher LSD post hoc test or with Student’s t-test unless otherwise noted in the figure legend. Values reported in the text and error bars are the mean ± SEM unless otherwise noted. All datasets were screened for outliers prior to analysis via Grubb’s test (α=0.05) and outliers were subsequently excluded. Statistical tests were performed in R or Prism 7 (GraphPad). Nonparametric tests were used where appropriate and tests were 2-tailed unless otherwise noted. For all experiments, *n* indicates the number of biological replicates. For cell culture experiments, this indicates the number of independently growing flasks or wells. For experiments involving animal behavior, this indicates the number of animals used. For experiments involving tissue collection from animals, this indicates the number of animals we collected the tissue from.

### GTEx data

Data from the GTEx Analysis Release V7 (dbGaP Accession phs000424.v7.p2) were obtained via the GTEx portal web tool. Expression values plotted are in transcripts per million (TPM), using the GENCODE annotated transcript for isoforms or a gene level model based on the GENCODE model with isoforms collapsed to single genes. Isoform expression values were hierarchically clustered using Euclidean distance and average linkage; dendrogram scale shows cluster distance.

### Analysis of bulk RNAseq and ChIPseq data

Single or paired-end RNAseq data was imported into the public Galaxy server at usegalaxy.org directly from the European Nucleotide Archive (study accession numbers PRJEB9006 and PRJNA262674) in FASTQ format and run through a standardized workflow consisting of quality trimming via Trim Galore!^39^ (Galaxy Version 0.4.2), read alignment to via HISAT^40^ (Galaxy Version 2.0.3), and feature counting via featureCounts (Galaxy Version 1.4.6.p5). Individual count files were grouped by treatment (Animal age) and differential expression testing was performed using DESeq2^41^ (Galaxy Version 2.11.39). All reference genomes and annotations were obtained from Gencode releases current at the time of analysis, including the Genome Reference Consortium Mouse Build 38 patch release 5 (GRCm38.p5) and evidence-based annotation of the mouse genome (GRCm38), version M16 (Ensembl 91), human build GRCh38 and the human annotation Release 25 (GRCh38.p7). Gene ontology (GO) enrichment was assessed using a PANTHER Overrepresentation Test web tool provided by the Gene Ontology Consortium^42,43^ (release date 2017-11-28). DAVID functional annotation was used to assess gene set enrichment for GAD_DISEASE_CLASS using default settings (DAVID 6.8).

CHART-seq data was accessed via NIH SRA Toolkit from accession PRJNA252626 and analyzed using similar read quality control and alignment tools as described above. CHART-seq peaks were called using the MACS2 algorithm^44,45^. Overlapping peaks were combined into a single peak, as recommended for input into ChIP-Enrich package. Using the ChIP-Enrich R package^46^ (version 2.4.0), CHART-seq peaks from MACS2 were assigned to the nearest transcription start site and GO Enrichment was assessed for Biological Processes and Molecular Functions.

### scRNA-seq analysis

Data were obtained via the European Bioinformatics Institute’s Single-cell Expression Atlas. T-distributed Stochastic Neighbor Embedding (t-SNE) plots were constructed using transcript per million (TPM) values from the transcriptomes of 3,589 single cells biopsied from four glioblastoma patients^22^. Unbiased clusters were generated using a t-SNE perplexity of 10; plots were colored via biased inferred cell type, as reported by the authors of the dataset. Biopsied tissue included cells from the tumor core as well as peripheral tissue; however, all cells inferred to be neurons were collected from noncancerous tissue adjacent to the glioblastoma.

## SUPPLEMENTAL FIGURES

**Supplemental Figure S1.**
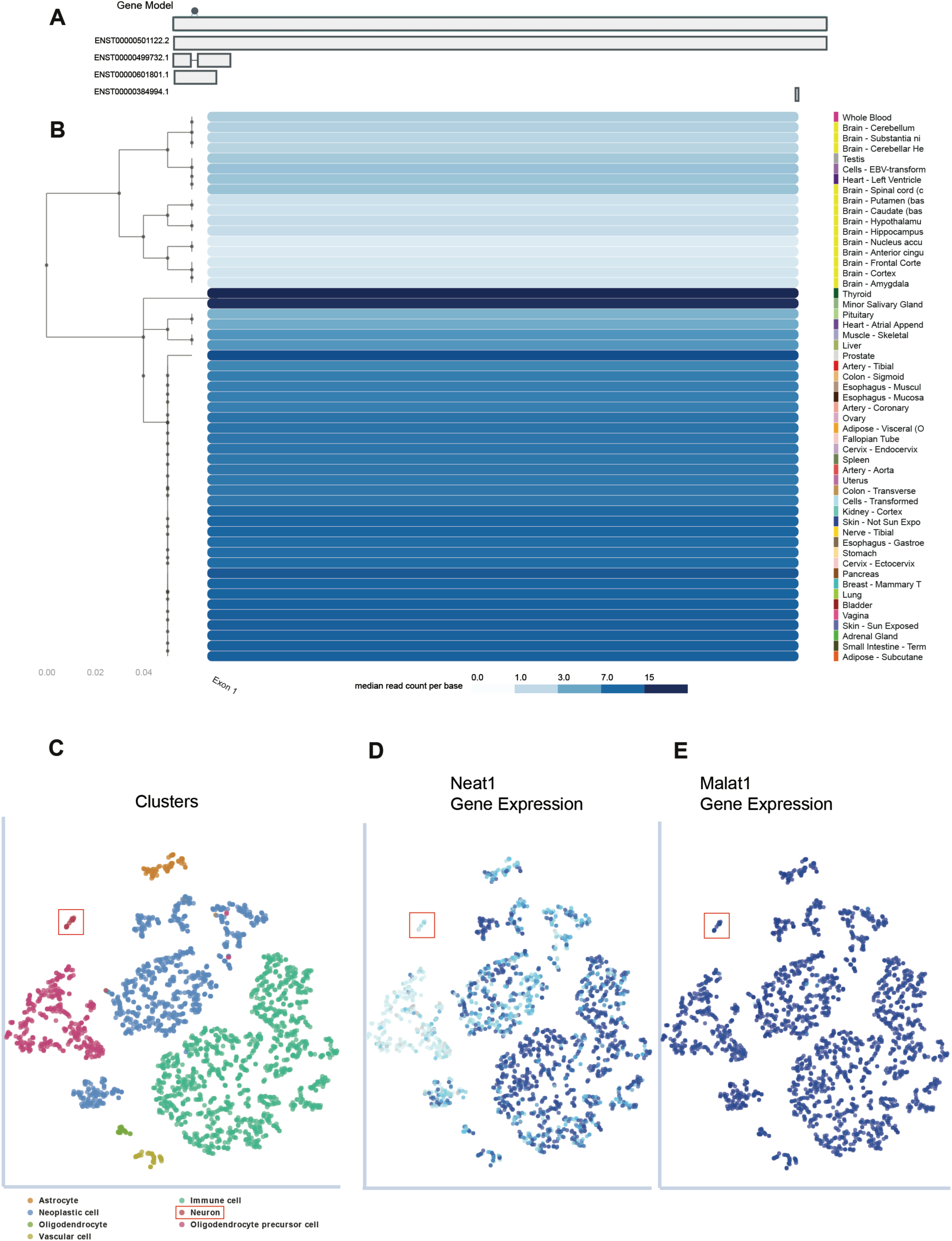
*NEAT1* expression is uniquely reduced in the human CNS, and baseline expression is low in human neurons relative to other cell types. **(A)** Diagram of *NEAT1* gene structure used in hierarchical clustering of *NEAT1* (B) Hierarchical clustering of *NEAT1* based on exon level expression in 53 human tissues from the GTEx project. Dendrogram scale shows cluster distance. Expression values displayed in the heatmap are the median expression values in TPM for each exon in each tissue. (C-E) t-SNE plots constructed using transcript per million (TPM) values from the transcriptomes of 3,589 biopsied human single cells. (C) Inferred cell types in unique colors, with clustered neurons outlined in red. (D) Expression of *NEAT1* in single cells heatmap, from 0 to 1200 TPM. (E) Expression of MALAT1 (NEAT2) in single cells, from 0 to 12000 TPM.

**Supplemental Figure S2.**
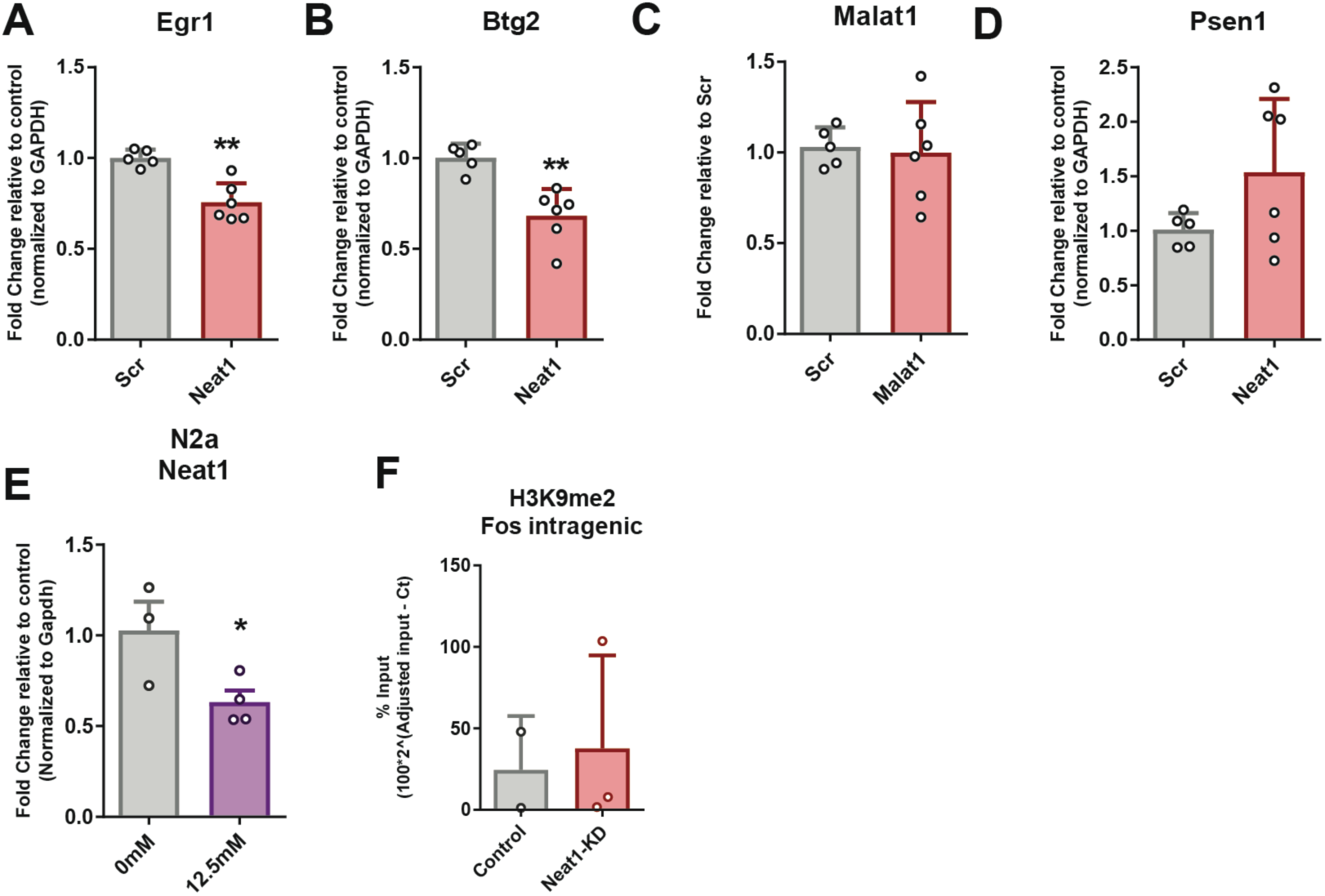
Neuronal regulation of immediate early genes after *Neat1* knockdown. (A-D) Differential expression of the immediate early genes (A) Egr1 (*n* = 5,6; *p* = 0.0010), (B) Btg2 (*n* = 5,6; *p* = 0.0018), (C) MALAT1 (*n* = 5,6; *p* = 0.8133), and (D) Psen1 (*n* = 5,6; *p* = 0.1228) after knockdown of *Neat1* in cultured neuronal cells. (E) Expression of the lncRNA *Neat1* in N2a cells after treatment with KCl (*n* = 3,4; *p* = 0.0495) (F) H3K9me2 at an intragenic region of the *c-Fos* gene is unchanged after *Neat1* knockdown in N2a cells (*n* = 2,3; *p* = 0.7939).

**Supplemental Figure S3.**
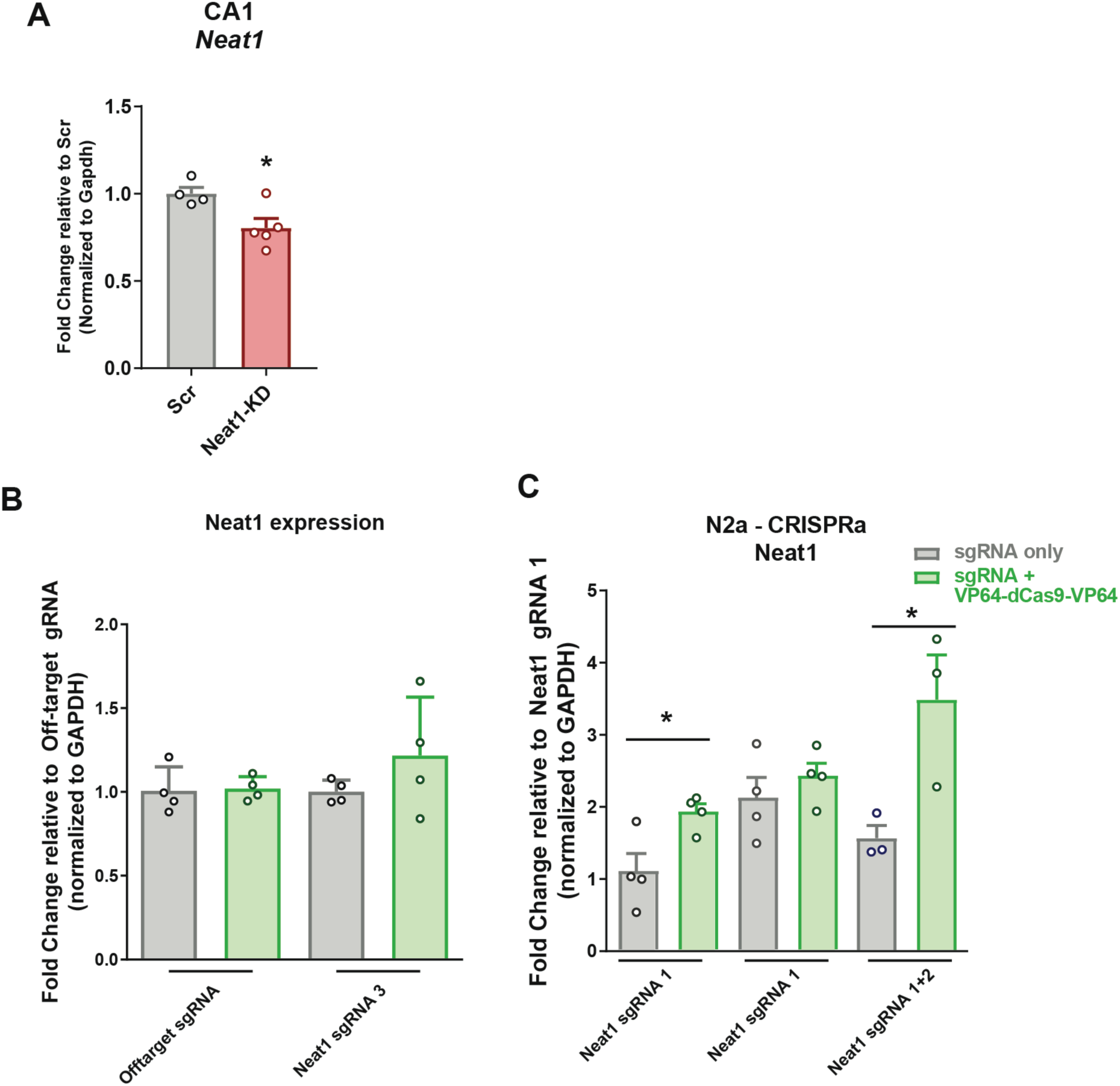
Validation of *Neat1* expression manipulation via RNAi and CRISPRa. (A) RT-qPCR quantification of *Neat1* expression in dCA1 of naïve mice 5d after *in vivo* transfection. (*n* = 4,5; *p* = 0.0242) (B-C) RT-qPCR quantification of *Neat1* in response to three different sgRNAs for CRISPRa-mediated upregulation of *Neat1.* (B) Transfection of an off-target sgRNA plasmid alone, *Neat1*-targeting sgRNA 3 plasmid alone, or off-target sgRNA plasmid and ddCas9-2xVP64, while sgRNA 3 in conjunction with dCas9-2xVP64 had a modest effect on expression (not statistically significant; *p* > 0.05). (C) Transfection of *Neat1* sgRNA plasmids #1 or #1/#2 in conjunction with dCas9-2xVP64 results in more robust transcriptional and additive upregulation of *Neat1* relative to cells receiving only the respective sgRNAs *(n* = 4,4,4,4,3,3; *\*p* < 0.05).

**Supplemental Figure S4.**
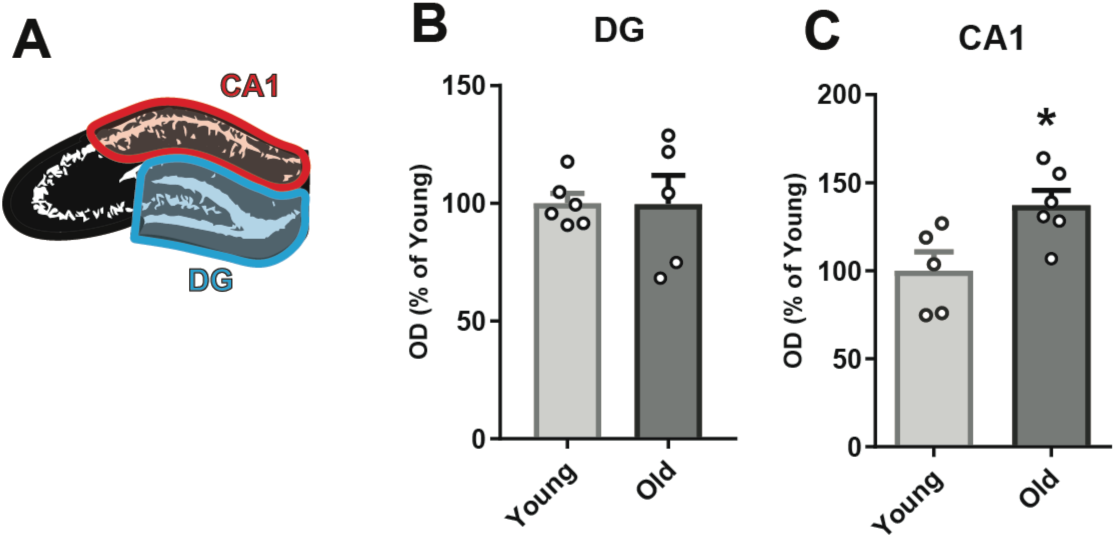
Age-related elevation of H3K9me2 in dCA1. (A) Cartoon of dissection of CA1 from DG in *Rattus norvegicus*. (B) DG expression of H3K9me2 is unchanged between young and aged rats (*n* = 6,6; *p* = 0.9762), while expression of H3K9me2 in CA1 (C) is elevated with aging (*n* = 5,6; *p* = 0.0465).

**Supplemental Figure S5.**
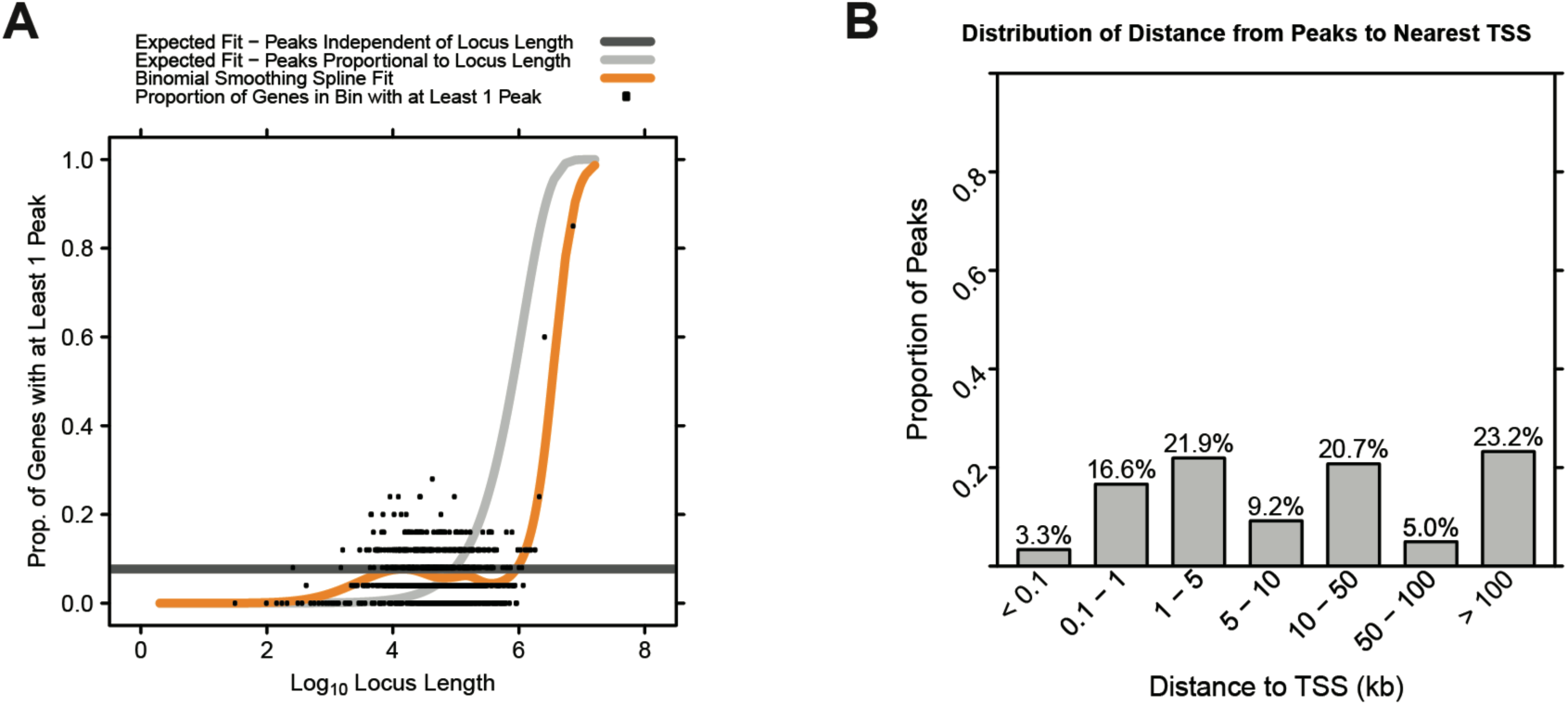
Quality control plots from Chip-Enrich. (A) Curves comparing *NEAT1-*bound peaks to locus length. (B) Distribution of distance from *NEAT1* bound peaks to the nearest gene TSS

